# Rad18 suppresses the late step of human immunodeficiency virus type 1 replication

**DOI:** 10.1101/2025.11.26.690725

**Authors:** Yasuo Ariumi, Chengcheng Zou, Hiroyuki Fukuda, Satoshi Tateishi, Eri Miyagi, Klaus Strebel

## Abstract

The Rad18 DNA repair protein plays a central role in the response to DNA damage and has been suggested to bind to the human immunodeficiency virus type 1 (HIV-1) integrase (IN) and restrict the early steps of HIV-1 infection. However, the inhibitory mechanism of Rad18 is not fully understood. Here, we describe a series of experiments that indicate that Rad18 can suppress the late step of HIV-1 replication by inhibiting selective viral post-transcription and the production of infectious HIV-1 virions. Notably, Rad18 interacted with HIV-1 IN, Tat, and Vif proteins and hijacked them in the nucleoli. Therefore, Rad18 seems to suppress multiple steps of HIV-1 infection, including viral post-transcription, production, and infectivity.

**Importance:** Rad18, which contributes to damage bypass and DNA post-replication repair, is known to bind to HIV-1 IN and suppress HIV-1 infection. In this study, we found for the first time that Rad18 suppresses the late stages of HIV-1 infection. In addition to IN, we identified Tat and Vif as Rad18-interacting proteins. Rad18 may suppress multiple steps of the HIV-1 life cycle, including the early steps of HIV-1 infection, reverse transcription and integration, and the late steps of HIV-1 infection, viral post-transcription, viral production, and viral infectivity. These findings highlight the protective role of DNA repair proteins against retroviral infection and L1 retrotransposition as guardians of the human genome.

## Introduction

DNA damage tolerance (DDT) pathways play a key role in allowing replication to proceed in the presence of DNA lesions in the DNA template. DNA lesions caused by ultraviolet (UV) radiation or chemical agents are removed and repaired by nucleotide excision repair (NER); however, DNA damage persists during DNA synthesis and causes DNA replication to stall. To avoid stalling, Rad18, a post-replication repair protein, contributes to damage bypass and DNA repair (1-3). Rad18 is an E3 ubiquitin ligase that is recruited to DNA damage sites. The DNA replication fork stalled by DNA damage induces mono-ubiquitination of proliferating cell nuclear antigen (PCNA), an essential replication accessory factor, on lysine (K)164 by the Rad6 E2 ubiquitin-conjugating enzyme/Rad18 E3 ubiquitin ligase complex (4-6). The bypass of damaged DNA occurs via two pathways: translesion synthesis (TLS) and template switching (TS). Mono-ubiquitination of PCNA promotes error-prone TLS, and Rad5-mediated poly-ubiquitination of PCNA on K63 promotes the TS pathway. PCNA mono-ubiquitination triggers the replacement of replicative DNA polymerase δ with translesion synthesis polymerase η, which can replicate DNA lesions (7-9). Rad18 deficiency causes hypersensitivity to various DNA-damaging agents, defects in post-replication repair, genomic instability, and an increase in sister chromatid exchange (10-12), suggesting that Rad18 plays an essential role in maintaining chromosomal DNA.

Furthermore, we recently found that Rad18 restricts long interspersed element-1 (LINE-1, L1) and Alu retrotransposition as a guardian of the human genome against endogenous retroelements (13). Rad18 interacts with human immunodeficiency virus type 1 (HIV-1) integrase (IN) (14), which catalyzes the HIV-1 integration process and suppresses the early steps of HIV-1 infection (15). However, the mechanism by which Rad18 suppresses HIV-1 infection is not fully understood. In this study, we further investigated the molecular mechanisms the anti-viral effect against HIV-1 and demonstrated a novel role for Rad18 in the late stages of HIV-1 replication.

## Results

### Rad18 suppresses late-stage HIV-1 replication

To study the potential role of Rad18 in HIV-1 infection, we first used lentiviral vector-mediated RNA interference to stably knockdown Rad18 in CD4^+^HeLa P4.2 cells, which constitutively express CD4, an HIV-1 receptor (16). We used puromycin-resistant pooled cells 10 days after lentiviral transduction. Western blotting showed a very effective knockdown of the endogenous Rad18 protein in P4.2 cells transduced with lentiviral vectors expressing shRNA targeted to Rad18 (Figure 1A). We found that the levels of the intracellular p24 capsid (CA) and its precursor form, pr55 Gag proteins, were significantly enhanced in Rad18 knockdown cells 48h after HIV-1 infection (Figure 1A), suggesting that Rad18 suppresses HIV-1 infection. Importantly, the shRNA did not affect cell viability (data not shown). We next analyzed the HIV-1 titer using a luciferase assay in the supernatants from the control (shCont) or Rad18 knockdown (shRad18) cells 48h post-infection with HIV-1 (strain of R9 or NL4-3) (17, 18). For this, we used CD4^+^ CXCR4^+^CCR5^+^HeLa TZM-bl reporter cells, in which both the luciferase and-galactosidase genes were separately integrated under the control of the HIV-1 promoter. The HIV-1 Tat protein, expressed as an early protein in HIV-1 infected cells can trans-activate the HIV-1 long terminal repeat (LTR) promoter, resulting in elevated luciferase activity. Culture supernatants (100 μl of culture supernatants were inoculated into TZM-bl cells, and luciferase activity 24h later. Consequently, the HIV-1 titer was enhanced (three-to four-fold) in the Rad18 knockdown cells compared with that in the control cells (Figure 1B), indicating that HIV-1 production was enhanced in the Rad18 knockdown cells. Thus, Rad18 may act as an anti-HIV-1 protein.

**Figure 1.**
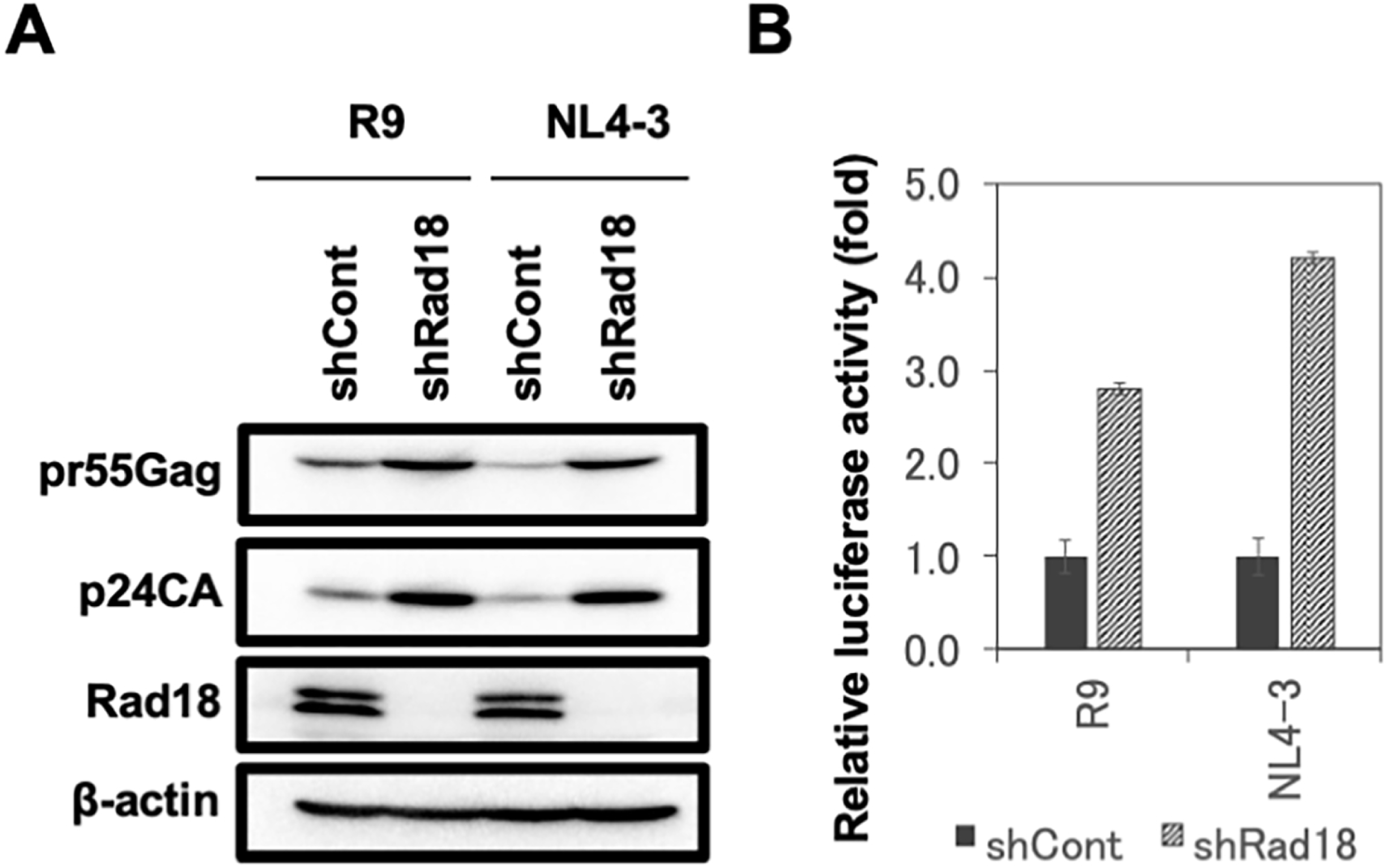
Rad18 suppresses HIV-1 infection. (**A**) HIV-1 p24 protein expression levels in Rad18 knockdown P4.2 cells. Inhibition of endogenous Rad18 protein expression by shRNA-producing lentiviral vector. The results of western blot analysis of cellular lysates with anti-Rad18 (ab188235 and ab79763; Abcam), anti-HIV-1 p24 (65-005; Bioacademia), or anti-β-actin antibody in HIV-1 (R9 or NL4-3)-infected P4.2 cells at 72 h post-infection are shown. (**B**) Comparison of HIV-1 titers in culture supernatants from the control (shCont) or Rad18 knockdown P4.2 cells (shRad18) as shown in Fig. 1A. Each 100 μl of the supernatants was inoculated into TZM-bl cells, and luciferase assays were performed 24 h post-infection. Luciferase activity in the Rad18 knockdown cells was calculated relative to that in the control cells transduced with a control lentiviral vector.

To test this hypothesis, we performed a series of Rad18 overexpression studies in 293T cells using four different HIV-1 molecular clones, including CXCR4 (X4)-tropic R9 and NL4-3, and CCR5 (R5)-tropic JR-FL (19) and AD8 (20). Ectopic expression of Myc-tagged Rad18 strongly suppressed the extracellular p24 CA protein expression of all four HIV-1 strains in their culture supernatants from 293T cells co-transfected with Rad18-Myc-expressing plasmid and each HIV-1 molecular clone (Figure 2A). In contrast, Rad18-Myc did not significantly affect so much the intracellular p24 CA and Vpr protein expression (Figure 2A). In addition, Rad18-Myc did not decrease the intracellular p17 matrix (MA), Nef, or Vif protein expression (Figure 2A). HIV-1 envelope (Env) glycoprotein is expressed as a precursor glycoprotein, gp160, which is cleaved by cellular proteases into the mature surface glycoprotein gp120 and transmembrane glycoprotein gp41. Notably, Rad18-Myc markedly decreased intracellular gp160, gp120, and gp41 Env protein expression (Figure 2A and 3). We then analyzed the HIV-1 titer using a luciferase assay in the supernatants from 293T cells co-transfected with the HIV-1 molecular clone (NL4-3) and/or Rad18-Myc-expressing plasmid. The indicated amounts of culture supernatants were inoculated into TZM-bl cells, and luciferase activity was measured in the cell lysates 24h later. Consequently, the HIV-1 titer was strongly suppressed in Rad18-Myc-expressing cells compared to that in control cells (Figure 2B). Furthermore, we analyzed HIV-1 infectivity after normalizing for HIV-1 p24 CA in the supernatants using p24 ELISA. The normalized supernatants (same amounts of HIV-1 p24 CA input: approximately 10 ng) were inoculated into TZM-bl cells, and luciferase activity 24h later. Even after normalization, HIV-1 infectivity was significantly suppressed in Rad18-Myc-expressing cells (Figure 2C). Moreover, we examined whether Rad18 is incorporated into the HIV-1 virion because HIV-1 integrase (IN) is incorporated into the virion. However, Rad18-Myc was not incorporated into HIV-1 virions (Figure 2D). Thus, Rad18 appears to suppress infectious HIV-1 production.

**Figure 2.**
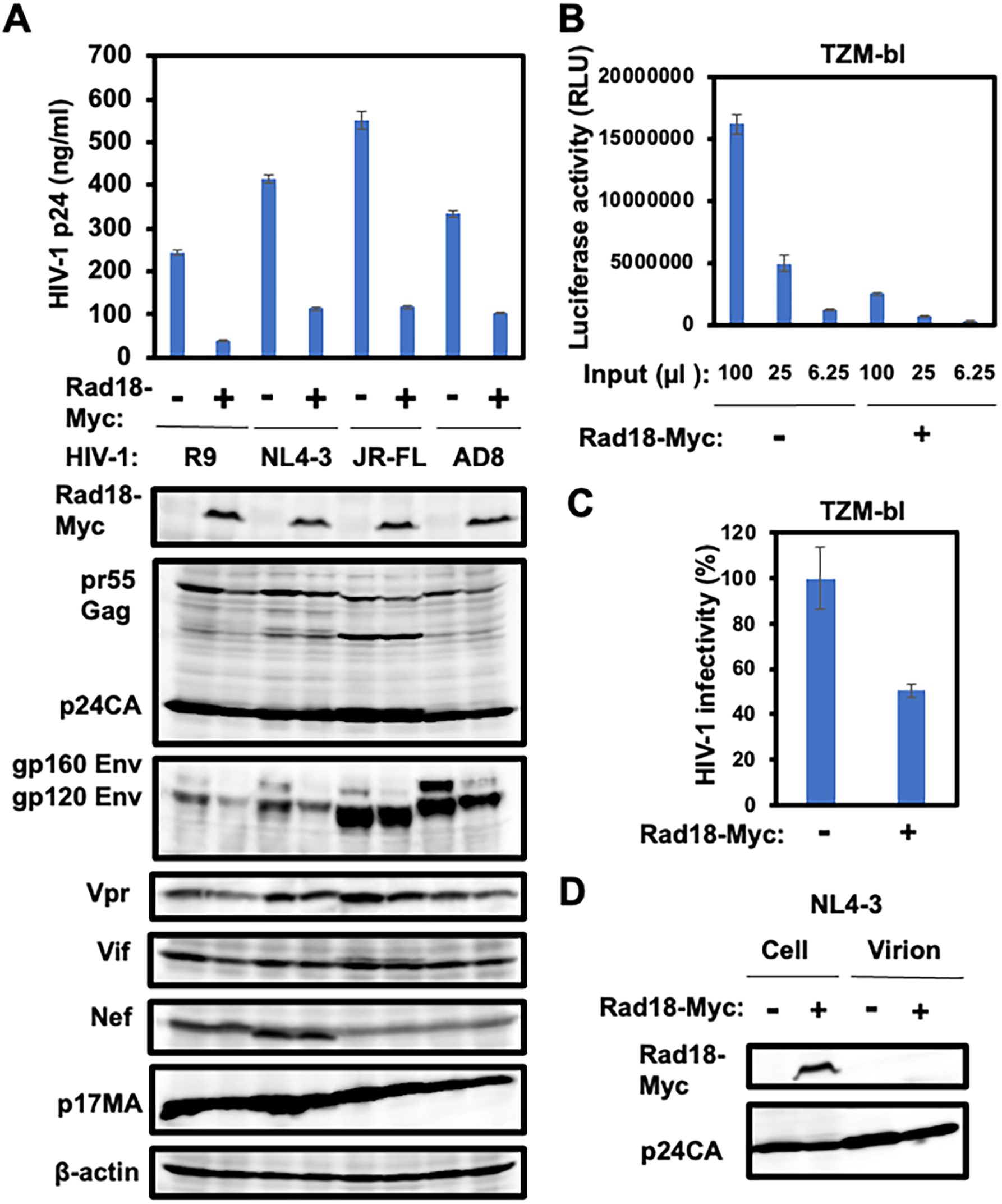
Rad18 overexpression suppresses HIV-1 production and viral infectivity. (**A**) 293T cells (2×10^5^ cells/well) were co-transfected with each HIV-1 molecular clone (2 μg) and pRad18-Myc or empty vector pCAG (2 μg). 48 h after transfection, p24 levels in the culture supernatants were measured using p24 ELISA. The results of Western blot analysis of cellular lysates with anti-Myc-tag (PL14; MBL), anti-HIV-1 p24 (65-005; Bioacademia), anti-HIV1 Vif (ab66643; Abcam), anti-HIV-1 GP41 (20-HR92; Fitzgerald) anti-HIV-1 Vpr (NCG-M01; Cosmo Bio), anti-HIV-1 Nef (3E6, Icosagen, Estonia), anti-HIV-1 p17 antibody (65-008, Bioacademia), or anti-β-actin antibody are shown. (**B**) Comparison of HIV-1 titer of culture supernatants from 293T cells (2×10^5^ cells/well) co-transfected with pNL4-3 (2 μg) and either pRad18-Myc or empty CAG vector (2 μg). 48 h after transfection, 100 μl, 25 μl, or 6.25 μl of the supernatants were inoculated into TZM-bl cells (2×10^4^ cells/well), and luciferase assays were performed 24 h post-infection. (C) HIV-1 infectivity after normalization with p24 ELISA. The normalized NL4-3 (same amount of p24 input) was inoculated into TZM-bl cells (2×10^4^ cells/well), and luciferase assays were performed 24 h post-infection. Luciferase activity in the presence of Rad18-Myc was calculated relative to that in the absence of Rad18-Myc. (D) Rad18 is not incorporated into the HIV-1 virion. 293T cells (2×10^5^ cells/well) were co-transfected with pNL4-3 (2 μg) and either pRad18-Myc or an empty CAG vector (2 μg). At 48 h post-transfection, the released HIV-1 virions were collected by centrifuging the culture supernatants from the transfected 293T cells at 20,000 × g for 2 h at 4 °C. The pellets containing virions were dissolved in lysis buffer and subjected to western blotting. The results of western blot analysis of cellular lysates or virions with anti-Myc-tag (PL14; MBL) or anti-HIV-1 p24 antibody (65-005; Bioacademia) are shown.

**Figure 3.**
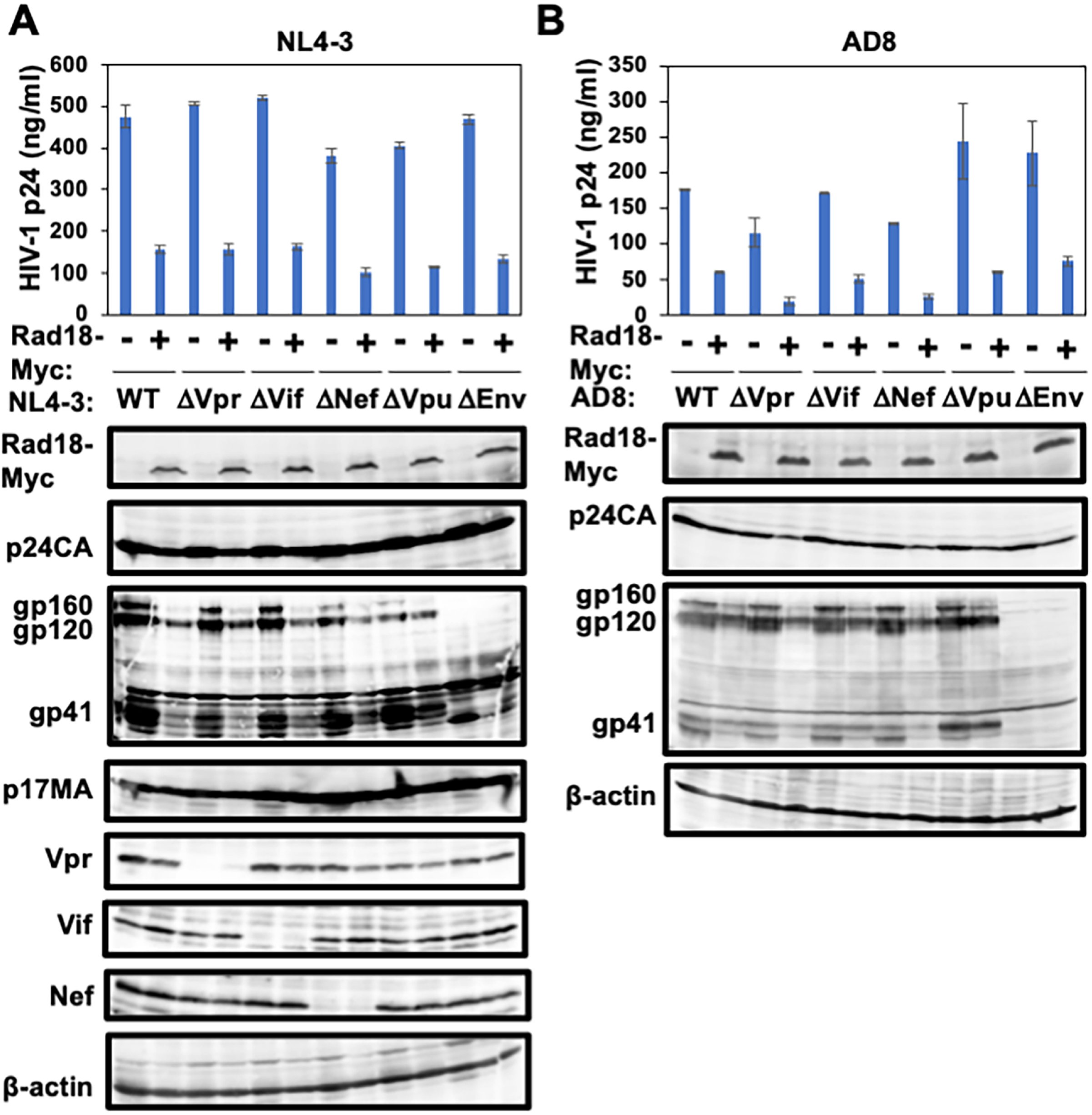
Effect of HIV-1 mutation on the antiviral activity of Rad18. 293T cells (2×10^5^ cells/well) were co-transfected with the HIV-1 molecular clones pNL4-3 (**A**) or pAD8 (**B**) (2 μg) and pRad18-Myc or the empty vector pCAG (2 μg). 48 h after transfection, p24 levels in the culture supernatants were measured using p24 ELISA. The results of western blot analysis of cellular lysates with anti-Myc-tag, anti-HIV-1 p24, anti-HIV-1 GP41, anti-HIV1 Vif, anti-HIV-1 Vpr, anti-HIV-1 Nef, anti-HIV-1 p17, or anti-β-actin antibodies are shown.

To identify the HIV-1 protein involved in the antiviral effect of Rad18, we used a wild-type (WT) HIV-1 molecular clone (NL4-3 or AD8) and deletion mutants of each viral protein (ΔVpr, ΔVif, ΔNef, ΔVpu, or ΔEnv) (20-26). Ectopic expression of Rad18 suppressed extracellular p24 CA protein expression in the culture supernatants and intracellular gp160/gp120 and gp41 Env protein expression in HIV-1 deletion mutants, similar to WT (Figure 3A and 3B). Notably, Rad18-Myc did not significantly affect so much the gp41 Env protein expression from HIV-1 ΔVpu (Figure 3A and 3B). Thus, these results suggest that at least four HIV-1 accessary proteins and Env are unrelated to the suppression of extracellular p24 CA expression by Rad18.

### Rad18 suppresses post-transcriptional step of HIV-1 replication

To further identify which HIV-1 replication step is involved in the antiviral effect of Rad18, we used pNL-A1, a derivative of pNL4-3 that lacks the *gag* and *pol* genes but expresses all other HIV-1 genes (27) (Figure 4A). Consequently, Rad18-Myc suppressed intracellular gp160/gp120 and gp41 Env as well as Vpr protein expression from pNL-A1 (Figure 4B), indicating that HIV-1 Gag and Pol proteins were dispensable for the inhibitory effect of Rad18. Importantly, Rad18-Myc did not affect Vif or Nef protein expression (Figure 4B), suggesting that Rad18 selectively suppresses the post-transcriptional step of HIV-1 replication. We obtained similar results using pNL-A1 (AD8), which replaced the NL4-3 Env with AD8 Env (Figure 4B). Unexpectedly, we noticed that vesicular stomatitis virus (VSV)-G Env reduced the level of Rad18-Myc protein expression, whereas Rad18-Myc did not affect VSV-G Env protein expression (Figure 4B). In contrast, Rad18-Myc downregulated JR-FL Env protein expression from the HIV-1 molecular clone JR-FL, whereas Rad18-Myc did not affect JR-FL Env protein expression only when JR-FL Env was co-expressed with Rad18-Myc (Figure 4C), indicating that Rad18 did not directly degrade HIV-1 Env. Furthermore, Rad18-Myc did not affect intracellular or extracellular HIV-1 p24 CA protein expression from cytomegalovirus (CMV) promoter-driven HIV-1 Gag and Pol packaging constructs pCMVΔR8.2 and pCMVΔR8.74 (28, 29) (Figure 4D). pCMVΔR8.2 expresses all HIV-1 proteins except Env, and pCMVΔR8.74 lacks four accessory proteins, Nef, Vif, Vpr, and Vpu, as well as Env (28, 29). Therefore, the LTR sequence may be critical for the inhibitory effect of Rad18.

**Figure 4.**
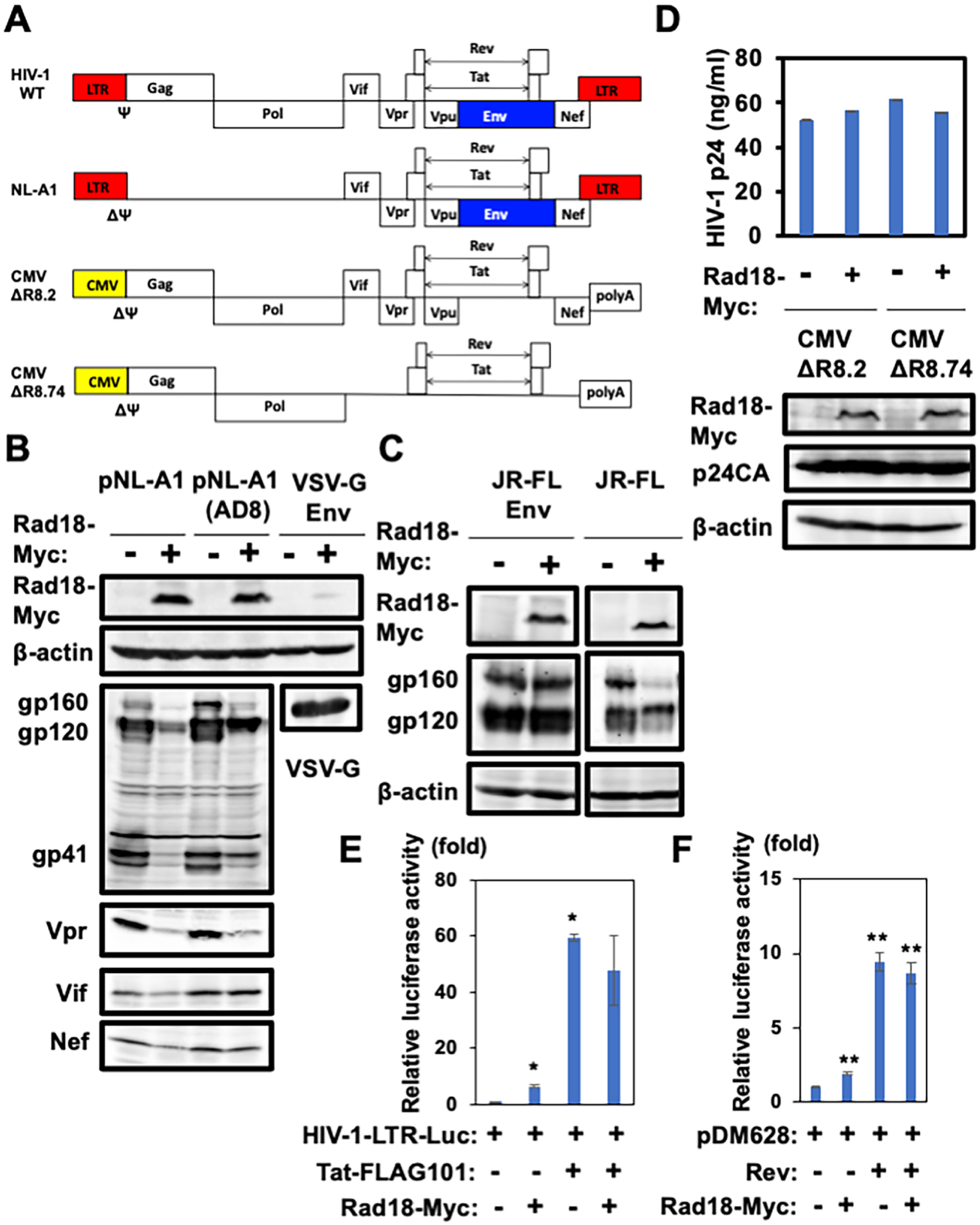
Selective post-transcriptional regulation of HIV-1 by Rad18. (**A**) Schematic representation of HIV-1 wild-type (NL4-3 WT), pNL-A1, pCMVΔR8.2, or pCMVΔR8.74 used in the present study. Ψ denotes the packaging signal. (**B**) 293T cells (2×10^5^ cells/well) were co-transfected with pNL-A1, pNL-A1 (AD8), or pVSV-G (2 μg) and pRad18-Myc or empty vector pCAG (2 μg). Cell lysates were prepared 48 h after transfection. The results of western blot analysis of cellular lysates with anti-Myc-tag, anti-HIV-1 GP41, anti-HIV1 Vif, anti-HIV-1 Vpr, anti-HIV-1 Nef, anti-VSV glycoprotein (P5D4; Sigma), or anti-β-actin antibody are shown. (**C**) 293T cells (2×10^5^ cells/well) were co-transfected with pJR-FL Env (65) or pJR-FL (2 μg) and pRad18-Myc or empty vector pCAG (2 μg). After 48 h of transfection, cell lysates were prepared. The results of western blot analysis of cellular lysates with anti-Myc-tag, anti-HIV-1 GP41, or anti-β-actin antibody are shown. (**D**) 293T cells (2×10^5^ cells/well) were co-transfected with pCMVΔR8.2 or pCMVΔR8.74 (2 µg) and pRad18-Myc or the empty vector pCAG (2 μg). 48 h after transfection, p24 levels in the culture supernatants were measured using p24 ELISA. The results of western blot analysis of cellular lysates with anti-Myc-tag (PL14; MBL), anti-HIV-1 p24 (65-005; Bioacademia), or anti-β-actin antibody are shown. (**E**) 293T cells (2×10^4^ cells/well) were co-transfected with pHIV-1-LTR-Luc (100 ng), pcDNA3-Tat101-FLAG (100 ng), and pRad18-Myc or the empty vector pCAG (100 ng), and luciferase assays were performed 24 h post-infection. The graph shows the mean (±SEM) firefly luciferase activity, with the condition without Rad18-Myc set to 100% (\**p*<0.05). (**F**) 293T cells (2×10^4^ cells/well) were co-transfected with pDM628 (100 ng), pcRev (100 ng), and pRad18-Myc or the empty vector pCAG (100 ng), and luciferase assays were performed 24 h post-infection. The graph shows the mean (±SEM) firefly luciferase activity with the condition without Rad18-Myc set to 100% (\*\**p*<0.01).

To confirm this possibility, we examined the impact of Rad18 on Tat-mediated HIV-1 transcription using a luciferase assay with the HIV-1-LTR-Luc reporter plasmid. 293T cells were co-transfected with the HIV-1-LTR-Luc reporter, Tat101-FLAG (30, 31), and/or Rad18-Myc plasmids, and a luciferase assay was performed 24 h later. Consequently, overexpression of Rad18 weakly inhibited Tat-mediated HIV-1 transcription; however, the impact of Rad18 on Tat-mediated transcription was not deemed significant by statistical analysis (Figure 4E). In addition, Rad18-Myc slightly enhanced the basal level of HIV-1 LTR promoter activity in the absence of Tat expression (Figure 4E). To further investigate the role of Rad18 in HIV-1 Rev function, we used a Rev-dependent luciferase-based reporter plasmid, pDM628 (32, 33). The luciferase level was markedly stimulated by Rev, which induced approximately a 10-fold increase in the reporter signal (Figure 4F). 293T cells were then co-transfected with Rad18-Myc and Rev expression plasmids. However, Rad18 had no effect on the luciferase level with Rev (Figure 4F), indicating that Rad18 does not regulate HIV-1 Rev function. Thus, HIV-1 transcriptional initiation and the Rev-dependent nuclear export step may not be critical targets, and Rad18 may regulate the HIV-1 post-transcriptional step.

Rad18 is an E3 ubiquitin ligase with multiple functional domains, including the N-terminal Really Interesting New Gene (RING)-finger domain (residues 25-63) commonly found in E3 ubiquitin ligases, required for interaction with the E2 ubiquitin ligase Rad6; the C2HC Zinc-finger DNA binding domain (residues 201-225), the SAF-A/B, Acinus, and Pias (SAP) DNA binding domain (residues 248-282), the C-terminal Rad6-binding domain (6BD) (residues 340-395), and the C-terminal Polη binding domain (C2) (residues 401-445) (Figure 5A) (4, 5). To determine which Rad18 functional domain is critical for the inhibitory activity against HIV-1, we used several Rad18 mutant-expressing plasmids (Figure 5A) (4, 13). An HIV-1 molecular clone NL4-3 and each Rad18 mutant-expressing plasmid were co-transfected, and we analyzed the extracellular p24 concentration by ELISA and the level of intracellular p24 CA protein expression by western blotting. Consequently, all Rad18 mutants suppressed extracellular p24 levels (Figure 5B). Nevertheless, the Rad18 mutants did not significantly affect so much the intracellular p24 CA expression (Figure 5B). Consistently, all Rad18 mutants suppressed intracellular gp160/gp120 and gp41 Env as well as Vpr protein expression from pNL-A1 (Figure 5C). These results suggest that Rad18 may regulate HIV-1 post-transcriptional steps and that Rad18 ubiquitination activity is independent of the inhibitory effect.

**Figure 5.**
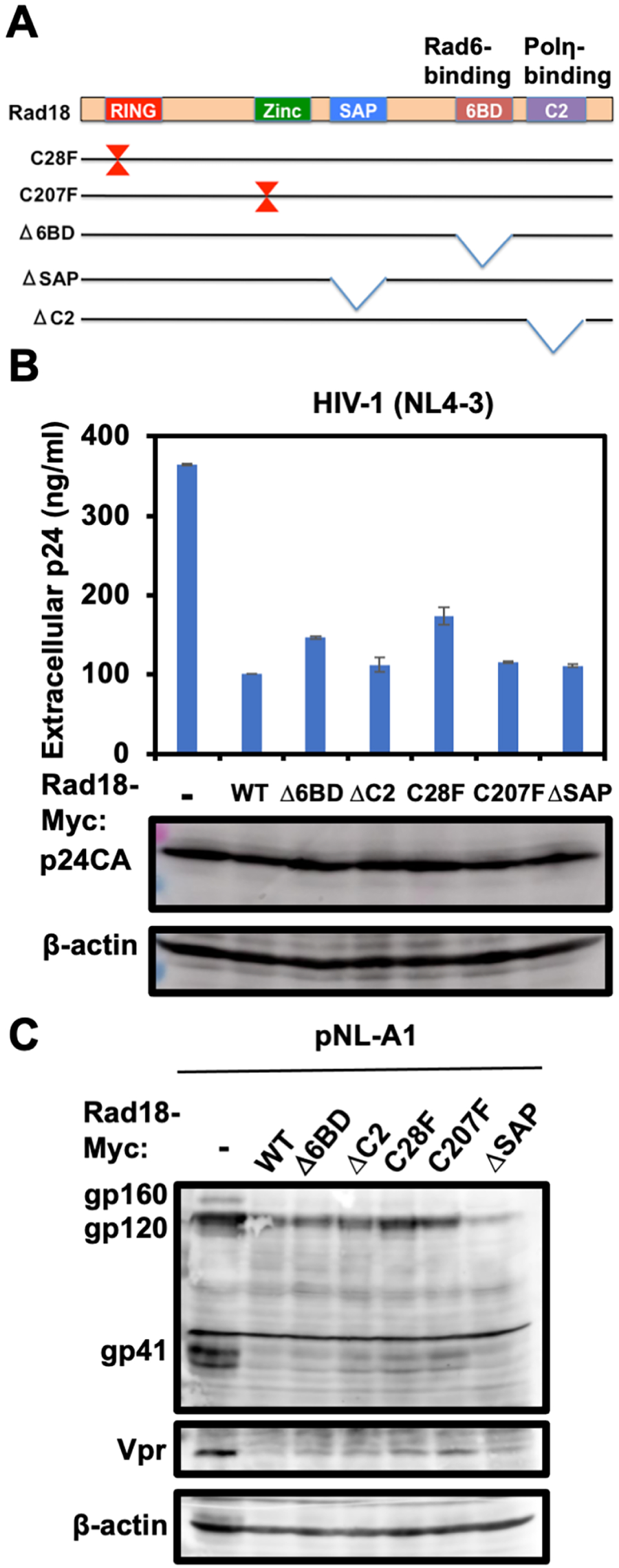
The ubiquitin ligase activity of Rad18 is dispensable for antiviral activity. (**A**) Schematic representation of Rad18 and the mutants used in this study. The RING finger, zinc finger, SAP, Rad6-binding (6BD), and C2 domains are indicated. (**B**) 293T cells were co-transfected with 2 µg of Myc-tagged Rad18 (WT, Δ6BD, ΔC2, C28F, C207F, or ΔSAP) expressing plasmid (4, 13) and 2 µg of pNL4-3. Protein expression levels of p24 CA and β-actin are shown by western blot analysis with anti-HIV-1 p24 or anti-β-actin antibody. (C) 293T cells were co-transfected with 2 µg of Myc-tagged Rad18 (WT, Δ6BD, ΔC2, C28F, C207F, or ΔSAP)-expressing plasmid (4, 13) and 2 µg of pNL-A1. Protein expression levels of gp160/gp120 Env, gp41 Env, Vpr, and β-actin are shown by western blot analysis with anti-HIV-1 GP41, anti-Vpr, or anti-β-actin antibodies.

### Rad18 is a nucleolar protein

To examine whether ectopically expressed Rad18 affects the subcellular localization of HIV-1 p24 CA or Env, HIV-1 molecular clone NL4-3 and Rad18-expressing plasmids were co-transfected into 293T cells. Consequently, HIV-1 p24 CA was predominantly localized in the plasma membrane and cytoplasmic punctate foci, and HIV-1 Env was predominantly localized in the perinuclear region (possibly Golgi apparatus) and plasma membrane (Figure 6). In contrast, ectopically expressed Rad18 formed a ring-like structure in the nucleus and did not affect the subcellular localization of the other proteins (Figure 6). Thus, HIV-1 p24 CA and Env are not direct targets of Rad18. However, little is known about the nuclear compartment in which Rad18 localizes.

**Figure 6.**
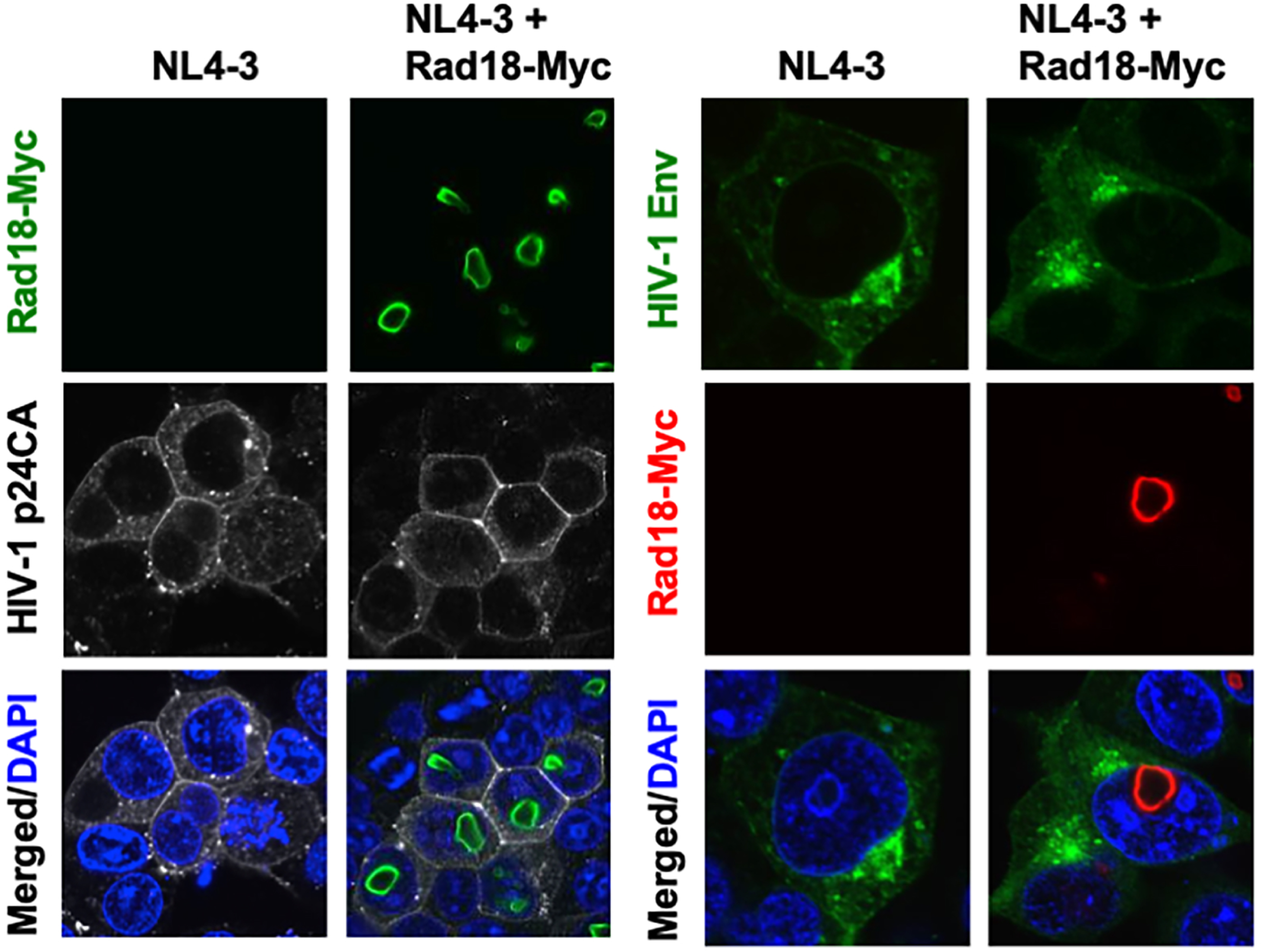
Subcellular localization of HIV-1 p24 capsid and Env in the presence or absence of ectopic Rad18 expression. 293T cells were transfected with 100 ng of the HIV-1 molecular clone NL4-3 or co-transfected with 100 ng of Myc-tagged Rad18. At 48 h post-transfection, the cells were stained with anti-HIV-1 p24 or anti-HIV-1 Env antibodies, and the signals were visualized using donkey anti-rabbit IgG antibody, Alexa Fluor 647, and anti-Myc-tag mAb-Alexa Fluor 488 (M047-A48; MBL), or goat anti-human IgG, Alexa Fluor 488, and anti-Myc-tag mAb-Alexa Fluor 594 (M047-A59; MBL).

To identify which nuclear compartment is the Rad18 localization site in the nucleus, we stained 293T cells using several nuclear body markers, anti-SC35 [nuclear speckles/interchromatin granule clusters (IGCs)], anti-Coilin (Cajal bodies), anti-PSF (Paraspeckles), anti-SMN (Gems), anti-PML [PML-nuclear bodies (NBs)], and anti-Nucleolin (Nucleoli), respectively. Consequently, endogenous Rad18 colocalized with nucleolin in the nucleolus, and Rad18 did not colocalize with other nuclear compartment markers (Figure 7A), indicating that Rad18 localizes to the nucleolus. Furthermore, we observed that endogenous Rad18 was juxtaposed with Coillin and PSF in the nucleolus (Figure 7A). Consistently, ectopically expressed Rad18 also colocalized with endogenous nucleolin (Figure 7B). Thus, we identified Rad18 as a nucleolar protein.

**Figure 7.**
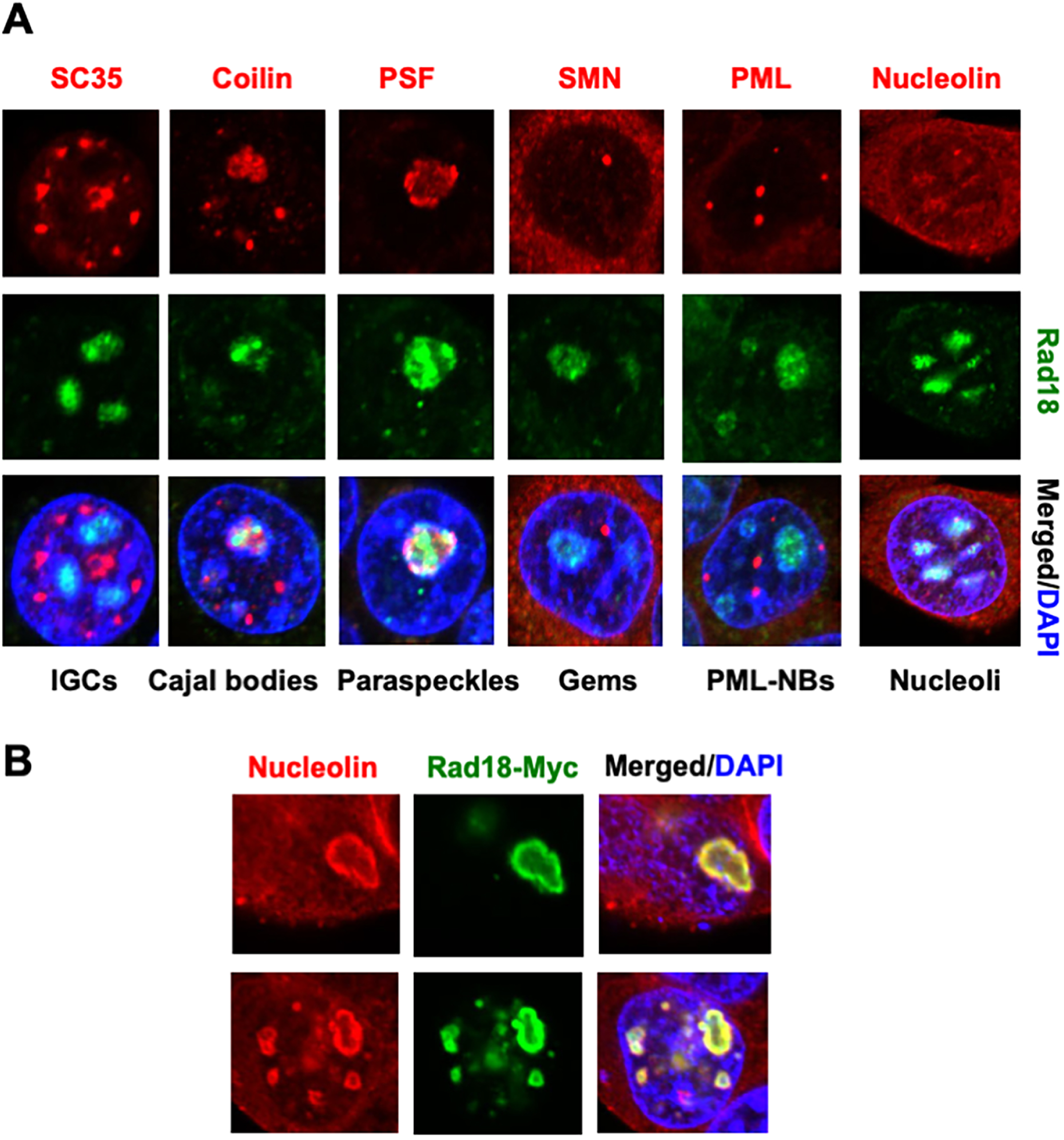
Rad18 localizes to the nucleoli. (**A**) Subcellular localization of endogenous Rad18 and various nuclear domain markers. 293T cells were stained with anti-SC-35 (SC-35, Sigma), anti-Coilin (ab11822; Abcam), anti-PML (PG-M3; Santa Cruz), anti-SFPQ (PSF) mAb (RN014MW; MBL), anti-SMN (610646; BD Transduction Lab), anti-Nucleolin mAb (4E2; MBL), and/or anti-RAD18 antibody (ab188235 and ab79763; Abcam). Cells were then stained with donkey anti-rabbit IgG, AlexaFluor488 conjugate, and donkey anti-mouse IgG secondary antibody, Alexa Fluor 594 conjugate. Images were visualized using a confocal laser scanning microscope. The nuclei were stained with DAPI (blue). (**B**) Ectopically expressed Rad-18 colocalizes with endogenous nucleolin. 293T cells were transfected with 100 ng of Myc-tagged Rad18. At 48 h post-transfection, the cells were stained with anti-nucleolin antibody (ab22758; Abcam), and the signals were visualized with anti-Myc-tag mAb-Alexa Fluor 488 (M047-A48; MBL) and donkey anti-mouse IgG Alexa Fluor 594.

### Rad18 interacts with HIV-1 integrase, Tat, and Vif

Rad18-Myc was localized in the Nucleolus and FLAG-tagged integrase (IN) was localized in the nuclear bodies (Figure 8A). When both proteins were co-expressed, Rad18-Myc hijacked and colocalized with FLAG-IN in Nucleolus (Figure 8B). In contrast, GFP-Vif colocalized with APOBEC3G (A3G)-HA, a cytidine deaminase with anti-HIV-1 activity, in cytoplasmic P-bodies in the absence of Rad18-Myc (Figure 8C). Notably, Rad18-Myc hijacked and colocalized with GFP-Vif from P-body to Nucleolus when both proteins were co-expressed (Figure 8D). Furthermore, HIV-1 Tat101-FLAG was localized in the nucleolus, nucleoplasm, and cytoplasm (Fig. 8E). Rad18-Myc predominantly colocalized with Tat101-FLAG in Nucleolus when both proteins were co-expressed (Figure 8E). In contrast, Rad18-Myc did not colocalize with HIV-1 Env (JR-FL strain) or Nef-FLAG (Figure 8F and 8G). Thus, Rad18 may interact with HIV-1 IN, Vif, and Tat.

**Figure 8.**
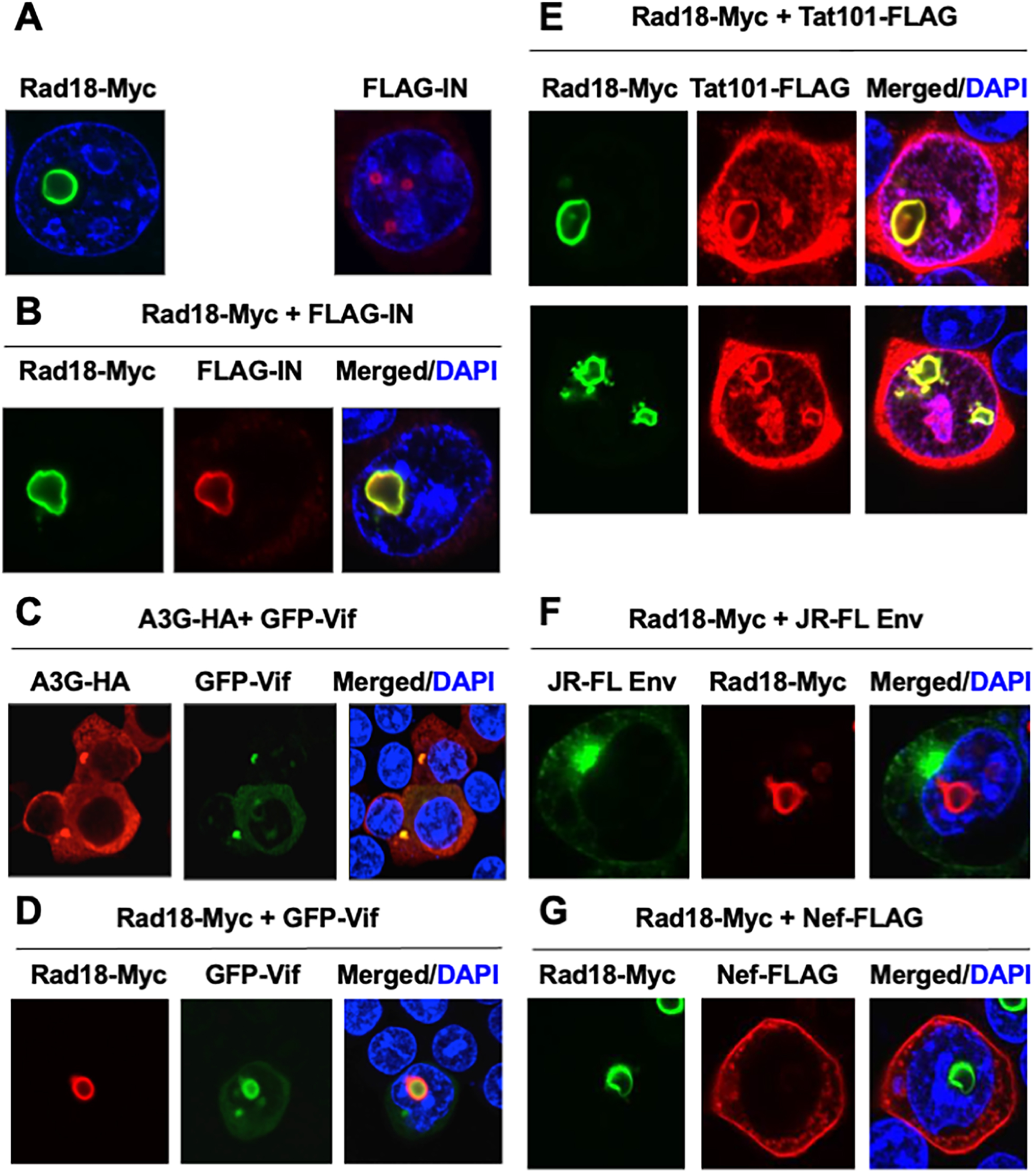
Rad18 colocalizes with HIV-1 integrase (IN), Vif, and Tat in the nucleoli. (**A**) Subcellular localization of Myc-tagged Rad18 or FLAG-tagged HIV-1 IN. 293T cells were transfected with 100 ng Myc-tagged Rad18 or 100 ng FLAG-tagged IN plasmid. At 48 h post-transfection, the cells were stained with anti-FLAG antibody (M2; Sigma-Aldrich), and the signals were visualized with anti-Myc-tag mAb-Alexa Fluor 488 or donkey anti-mouse IgG secondary antibody, Alexa Fluor 594. (**B**) Colocalization of Rad18 with HIV-1 IN when 293T cells were co-transfected with 100 ng of Myc-tagged Rad18 and 100 ng of FLAG-tagged IN. (**C**) Colocalization of Rad18 with HIV-1 Tat when 293T cells were co-transfected with 100 ng of Myc-tagged Rad18 and 100 ng of FLAG-tagged Tat101. (**D**) Colocalization of HA-tagged A3G with GFP-Vif when 100 ng of pA3G-HA and 100 ng of pGFP-Vif were co-transfected. (**E**) Hijacking of GFP-Vif from the cytoplasmic P-body to the nucleolus by Rad18-Myc when 293T cells were co-transfected with 100 ng of pRad18-Myc and 100 ng of pGFP-Vif. (**F**) HIV-1 Env did not colocalize with Myc-tagged Rad18. 293T cells were transfected with 100 ng pRad18-Myc and 100 ng pJR-FL Env (65). (**G**) HIV-1 Nef did not colocalize with Myc-tagged Rad18. 293T cells were transfected with 100 ng pRad18-Myc and 100 ng pNef-FLAG.

To further confirm the possibility that Rad18 interacts with HIV-1 IN, Tat, or Vif, we performed immunoprecipitation analysis. Consistent with the immunofluorescence data, Rad18-Myc co-immunoprecipitated with FLAG-IN, Tat101-FLAG, and GFP-Vif (Figure 9A-C). In contrast, Rad18-Myc failed to bind gp160/gp120 Env, Rev, or Nef-FLAG (Figure 9D). Thus, Rad18 interacts with HIV-1 IN, Tat, and Vif.

**Figure 9.**
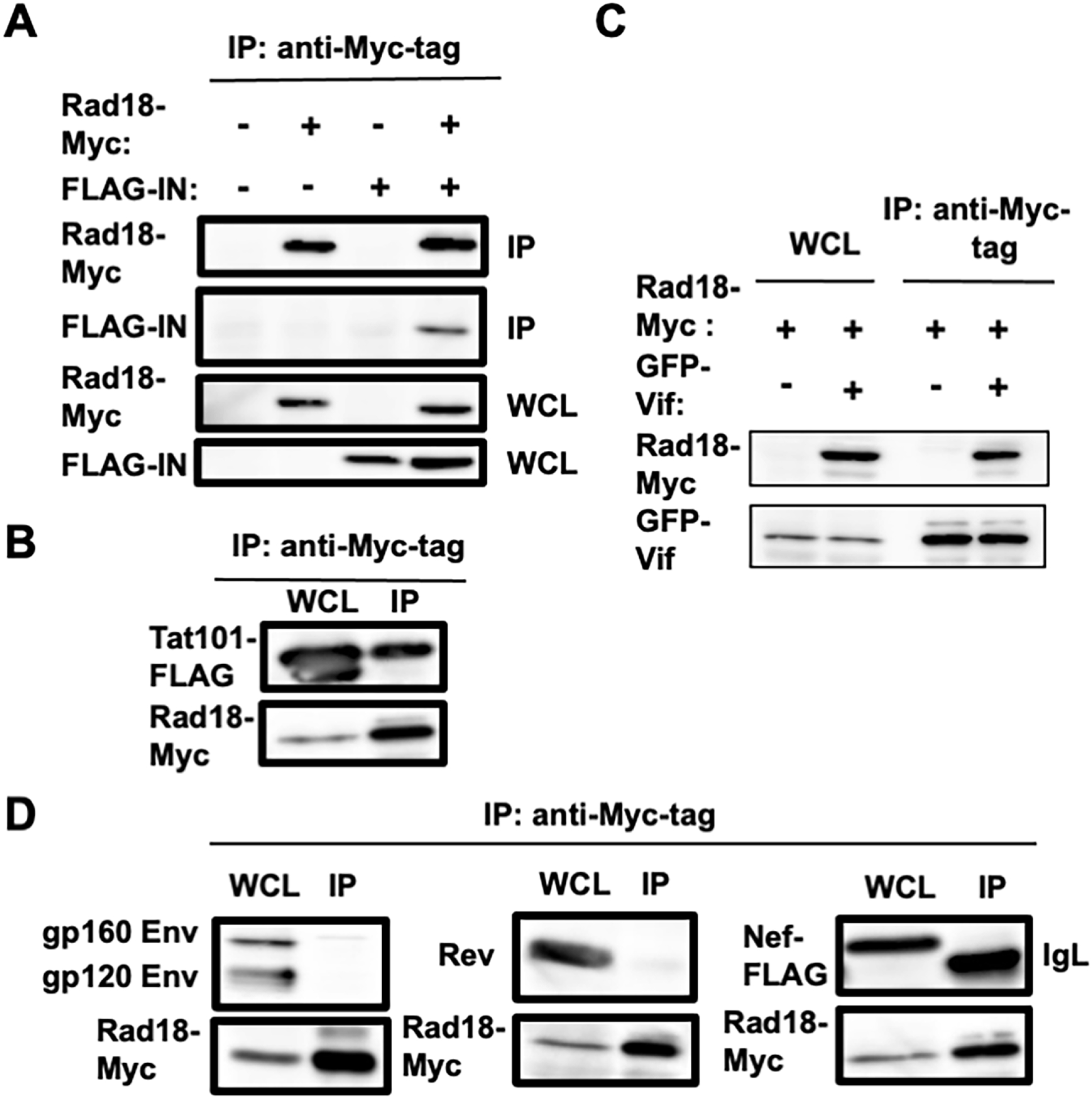
Rad18 binds to HIV-1 IN, Vif, and Tat. (A, B) 293T cells (2×105 cells/well) were co-transfected with 2 µg pRad18-Myc and 2 µg either pFLAG-IN (**A**) or pcDNA3-Tat101-FLAG (**B**). The cell lysates were immunoprecipitated with anti-Myc-tag antibody, followed by immunoblotting analysis using anti-Myc-tag or anti-FLAG antibody. The results of western blot analysis of 1/10 of the cellular lysates with anti-Myc-tag or anti-FLAG antibody are also shown. (**C**) 293T cells (2×10^5^ cells/well) were co-transfected with 2 µg pRad18-Myc and 2 µg pGFP-Vif. The cell lysates were immunoprecipitated with anti-Myc-tag antibody, followed by immunoblotting analysis using either anti-Myc-tag or anti-Living Colors A.v. monoclonal antibodies (JL-8; Clontech). (**D**) 293T cells (2×10^5^ cells/well) were co-transfected with 2 µg pRad18-Myc and 2 µg pJR-FL Env, pcRev, or pNef-FLAG. The cell lysates were immunoprecipitated with anti-Myc-tag antibody, followed by immunoblotting with anti-Myc-tag, anti-KD-247, or anti-FLAG antibody. IP: immunoprecip. WCL: whole cell lysate. IgL: immunoglobulin light chain.

To determine which Rad18 functional domain is required for binding to HIV-1 IN, Tat101-FLAG, or GFP-Vif, we used several Rad18 mutant-expressing plasmids (4, 6). Lysates of 293T cells co-expressing Myc-tagged Rad18 (WT, Δ6BD, ΔC2, C28F, C207F, or ΔSAP) and FLAG-IN, Tat101-FLAG, or GFP-Vif were immunoprecipitated with anti-Myc-tag antibody, followed by western blotting with anti-Myc-tag, anti-FLAG, or anti-GFP antibody. Consequently, FLAG-IN co-immunoprecipitated with WT Rad18, ΔC2, and C28F mutants (Figure 10A), and Tat101-FLAG co- immunoprecipitated with WT Rad18, ΔC2, C28F, C207F, and ΔSAP mutants (Figure 10B). However, the Δ6BD mutant failed to bind either FLAG-IN or Tat101-FLAG (Figure 10A and 10B), indicating that the Rad6-binding domain is critical for binding to HIV-1 IN or Tat. In contrast, GFP-Vif co-immunoprecipitated with WT Rad18 and the Δ6BD mutant (Figure 10C).

**Figure 10.**
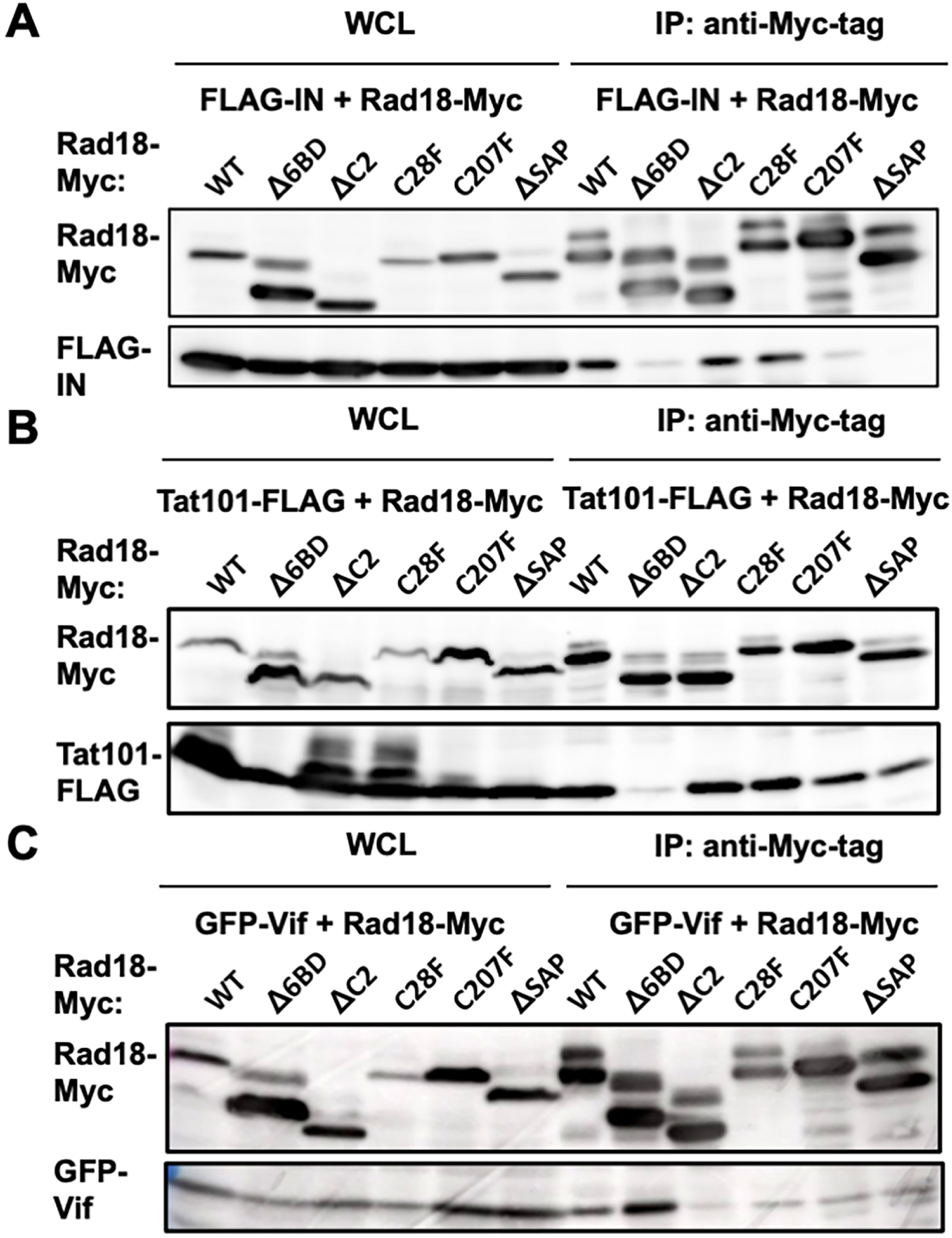
Rad18 functional domain required for interaction with HIV-1 IN, Tat, or Vif. 293T cells (2×10^6^ cells in a 10 cm dish) were co-transfected with 15 μg of pRad18-Myc (WT, Δ6BD, ΔC2, C28F, C207F, or ΔSAP) and 10 μg of pFLAG-IN (**A**), pcDNA3-Tat101-FLAG (**B**), or pGFP-Vif (**C**). The cell lysates were immunoprecipitated with anti-Myc-tag antibody, followed by immunoblotting analysis using anti-Myc-tag, anti-FLAG, and anti-Living Colors A.v. monoclonal antibody (JL-8; Clontech), respectively. WCL: whole cell lysates. IP: immunoprecipitation.

Finally, we examined whether HIV-1 IN, Tat, or Vif affected the ubiquitin ligase activity of Rad18. Rad18 is known to mono-ubiquitinate PCNA (4, 6). Rad18 could mono-ubiquitinate PCNA when Rad18-Myc, HA-Rad6B, HA-PCNA, and HA-Ub were co-expressed (Figure 11A). However, HIV-1 IN, Tat101, Vif, and Vpr did not suppress the mono-ubiquitination of PCNA by Rad18 (Figure 11B), suggesting that HIV-1 IN, Tat101, and Vif do not suppress the ubiquitin ligase activity of Rad18.

**Figure 11.**
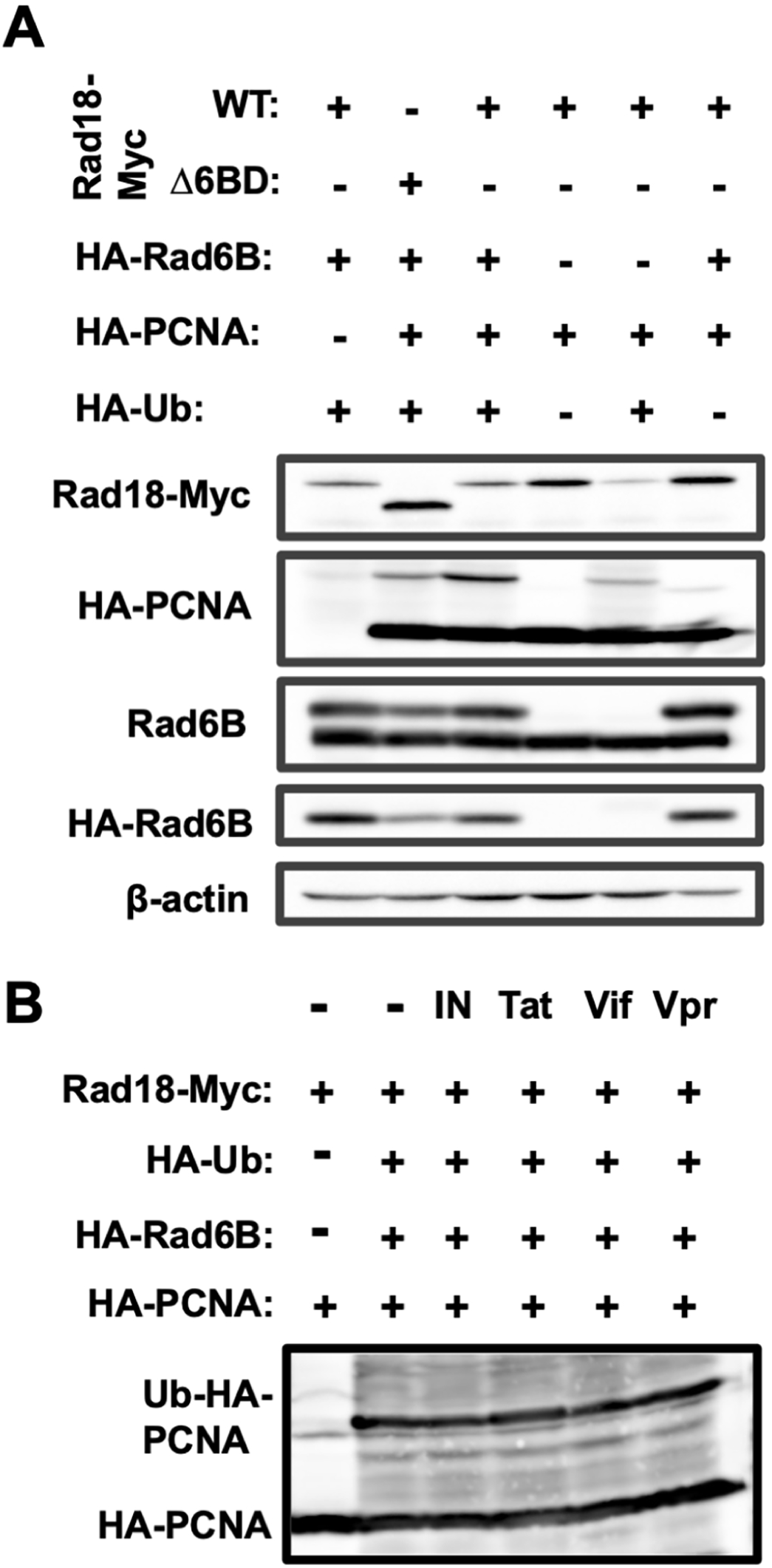
Mono-ubiquitination of PCNA by Rad18 is not inhibited by HIV-1 IN, Tat, Vif, or Vpr. (**A**) Mono-ubiquitination of PCNA by Rad18. 293T cells (2×10^5^ cells/well) were co-transfected with 1 μg of pRad18-Myc (WT or Δ6BD), pcDNA3-HA-Rad6B (13), pcDNA3-HA-PCNA and/or pHA-Ub (13). The results of western blot analysis of cellular lysates or virions with anti-Myc-tag, anti-HA (3F10), anti-Rad6B, or anti-β-actin antibodies are shown. (**B**) 293T cells (2×10^5^ cells/well) were co-transfected with 1 μg of pRad18-Myc, pcDNA3-HA-Rad6B, pcDNA3-HA-PCNA, pHA-Ub and/or pFLAG-IN, pcDNA3-Tat101-FLAG, pGFP-Vif, or pcDNA3-HA-Vpr. The results of western blot analysis of cellular lysates or virions with anti-HA antibody are shown.

## Discussion

Viral DNA integration is an essential step in the life cycle of HIV-1. HIV-1 integrase (IN) is a key player in this process. The initial genomic DNA cutting and joining steps of the integration process are mediated by HIV-1 IN; however, the final repair of residual DNA gaps must be repaired by host DNA repair enzymes, including cellular DNA polymerase (ß or δ) and DNA ligase (I, III, or IV and the cofactor XRCC4) (34). Unintegrated viral cDNA promotes apoptosis in non-homologous DNA end joining (NHEJ) component (Ku70, Ku80, DNA-PKcs, XRCC4, and DNA ligase IV) deficient cells (35). The NHEJ system circularizes unintegrated viral cDNA by end ligation and forms 2-LTR circles in infected cells to protect against apoptosis (35). Furthermore, several DNA damage sensors, such as ataxia-telangiectasia mutated (ATM), ataxia-telangiectasia and Rad3-related (ATR), DNA-dependent protein kinase (DNA-PK), and poly(ADP-ribose) polymerase-1 (PARP-1), play a central role in responses to various forms of DNA damage and have been suggested to facilitate HIV-1 integration and replication (36-46). Moreover, host cellular DNA repair proteins have been identified as HIV-1 IN-interacting proteins and are involved in HIV-1 integration. Ku70/Ku80 is the DNA-binding subunit of DNA-PK and plays a key role in NHEJ DNA repair. Ku70 interacts with HIV-1 IN and is involved in post-integration gap repair (47, 48). HIV-1 infection activates the Fanconi anemia (FA) DNA repair pathway, and HIV-1 IN interacts with the FA effector protein FANCI-D2 complex (49). Depletion of FANCI inhibits HIV-1 integration, suggesting that the FA pathway is required for HIV-1 integration (49).

In contrast, a protective role for DNA repair proteins against retroviral infection and L1 retrotransposition has been reported. Rad52, a component of homologous recombination (HR) DNA repair, suppresses HIV-1 infection (50). Rad51, another component of HR DNA repair, interacts with HIV-1 IN and inhibits its activity (51-53), whereas Rad51 activates HIV-1 LTR-dependent transcription to stimulate viral gene expression (54-56). In contrast, Ku represses HIV-1 transcription (57). Furthermore, Rad18, an E3 ubiquitin ligase, contributes to damage bypass, DNA post-replication repair, and HR DNA repair (1-3). Rad18 binds to HIV-1 IN and suppresses HIV-1 infection (14, 15). However, Lloyd *et al*. demonstrated integrase-catalytic activity-independent inhibition of HIV-1 by Rad18 (15). Rad18 deficient cells showed two-fold higher HIV-1 infectivity and increased reverse transcription products at earlier time points, suggesting that Rad18 suppresses reverse transcription (58). The DNA-binding-impaired mutant of Rad18 (L274P) failed to suppress HIV-1 infection, indicating that the DNA-binding activity of Rad18 is essential for suppressing HIV-1 infection (58). Rad18 binds to single-stranded DNA but not to double-stranded DNA (59), suggesting that HIV-1 suppression by Rad18 does not occur at the integration step. Therefore, Rad18 may inhibit reverse transcription by directly binding to the viral DNA intermediates. However, the details of the mechanism underlying the inhibitory effect of Rad18 are not fully understood. In this study, we further demonstrated that Rad18 suppresses the late stages of HIV-1 replication. Interestingly, Rad18 could suppress Vpr and Env expression levels but not Vif and Nef expression levels from pNL-A1 (Figure 4B) and suppresses the infectious virion production (Figure 2), suggesting that Rad18 affects the HIV-1 splicing pattern and that HIV-1 Gag and Pol proteins are dispensable for the inhibitory effect of Rad18. In this regard, Rad18 did not significantly affect so much for Tat-dependent HIV-1 transcription and Rev-dependent nuclear export function (Figure 4E and 4F). Therefore, Rad18 may selectively suppress post-transcription. Jacquenet *et al*. reported that the overexpression of SR proteins, including ASF/SF2, SC35, and 9G8, affects the HIV-1 splicing pattern, resulting in a drastic decrease in virus production (60). SC35 and 9G8 increased the levels of Tat RNA, whereas ASF/SF2 increased the levels of Vpr RNA. Importantly, all SR proteins examined negatively impacted the incorporation of Env TMgp41 in virions, again in agreement with the fact that Env mRNA and protein levels were drastically reduced in cells (60). Therefore, SR proteins can downregulate the late stages of HIV-1 replication. Rad18 may interact with and affect SR protein-mediated alternative HIV-1 RNA splicing patterns and virion production. These findings indicate that Rad18 plays different roles in the HIV-1 life cycle. We found that Rad18 interacts with HIV-1 IN, Tat, and Vif (Figure 8-10). Notably. Rad18 hijacked HIV-1 IN, Tat, and Vif into nucleoli (Figure 8). However, these interactions did not affect PCNA mono-ubiquitination by Rad18 (Figure 11). In contrast, Rad18 interacts with the Epstein-Barr virus (EBV) deubiquitinating enzyme, BPLF1, and contributes to the production of infectious viruses (61).

Furthermore, Rad18 restricts L1 and Alu retrotransposition as a guardian of the human genome against endogenous retroelements (13). Moreover, the nucleotide excision repair (NER) pathway, including XPA, XPC, XPD, XPF, and ERCC1, limits L1 retrotransposition (62, 63). DNA repair proteins act as components of broad innate immunity against incoming retroviruses and retrotransposition of endogenous retroelements. Indeed, APOBEC3G targets nascent retroviral cDNA during reverse transcription, whereas Rad18, Rad52, XPB, and XPD may target linear retroviral cDNA (64). Altogether, Rad18 seems to restrict HIV-1 infection at multiple steps of the HIV-1 life cycle. Rad18 may suppress multiple steps of the HIV-1 life cycle, including the early steps of HIV-1 infection, reverse transcription and integration, and the late steps of HIV-1 infection, viral post-transcription, viral production, and viral infectivity.

## Materials and Methods

### Cell culture

293T, P4.2 (16), and TZM-bl (NIH AIDS Research & Reference Program) cells were cultured in Dulbecco’s modified Eagle’s medium (DMEM; Life Technology, Carlsbad, CA, USA) supplemented with 10% fetal bovine serum (FBS). Transient transfection was performed using 293T cells and TransIT-LT1 (Mirus) according to the manufacturer’s protocol.

### Plasmid construction

To construct pcDNA3-HA-PCNA, a DNA fragment encoding PCNA was amplified from HuH-7 cDNA by PCR using KOD-Plus DNA polymerase (TOYOBO, Osaka, Japan) and the following pairs of primers: PCNA: 5’-CGGGATCCATGTTCGAGGCGCGCCTGGTCC-3’ (Forward), 5’-CCGGCGGCCGCCTAAGATCCTTCTTCATCC-3’ (Reverse). The obtained DNA fragments were subcloned into the *Bam*HI-*Not*I sites of the pcDNA3-HA vector, and the nucleotide sequences were determined by Sanger sequencing.

### HIV-1 molecular clones and Env expression plasmids

The different HIV-1 molecular clones used for transfection were pJR-FL (19), pAD8 (20), pNL4-3 (17), and pR9 (18). We also used a wild-type (WT) HIV-1 molecular clone (NL4-3 or AD8) and its mutants lacking each viral protein (ΔVpr, ΔVif, ΔNef, ΔVpu, or ΔEnv) (20-26). In addition, the HIV-1 JR-FL Env expression plasmid pJR-FL Env (65) or VSV-G Env expression plasmid pMD.2G (28, 29) were used.

### Collection and analysis of released virions

Released HIV-1 virions were collected by centrifuging the culture supernatants from transfected 293T cells at 20,000 × g for 2 h at 4 °C. The pellets containing virions were dissolved in lysis buffer and subjected to western blot analysis.

### RNA interference and lentiviral vector production

The vesicular stomatitis virus (VSV)-G-pseudotyped HIV-1-based vector system has been described previously (28, 29). Lentiviral vector particles were produced by transient transfection of 293T cells with the second-generation packaging construct pCMVΔR8.74 (28, 29) and the VSV-G-envelope-expressing plasmid pMD2G, as well as pLVshRad18 (13) or pLVshCon (13) lentiviral vectors into 293T cells with TransIT-LT1 transfection reagent (Mirus Bio LLC, Madison, WI, USA).

### Western blot analysis

Cells were lysed in a buffer containing 50 mM Tris-HCl (pH 8.0), 150 mM NaCl, 4 mM EDTA, 1% Nonidet (N) P-40, 0.1% sodium dodecyl sulfate (SDS), 1 mM dithiothreitol (DTT), and 1 mM phenyl-methylsulfonyl fluoride (PMSF). Supernatants from these lysates were subjected to SDS-polyacrylamide gel electrophoresis and transferred to Immobilon-P PVDF transfer membranes. The membranes were then incubated with the following primary antibodies: anti-HIV-1 p24 rabbit polyclonal antibody (65-004; Bioacademia, Japan), anti-HIV-1 Env (KD-247) (66), HIV1 GP41 antibody (20-HR92; Fitzgerald), Mouse monoclonal antibody to HIV-1 Nef (3E6, Icosagen, Estonia), Mouse monoclonal antibody HIV-1 Rev (10, Icosagen), anti-HIV-1 p17 antibody (65-008, Bioacademia), anti-HIV1 Vif antibody [319] (ab66643; Abcam), anti Vpr (HIV-1) monoclonal antibody (NCG-M01; Cosmo Bio, Japan), anti-Myc-tag antibody (PL14; MBL), anti-VSV Glycoprotein (P5D4; Sigma), anti-HA (HA-7; Sigma), and anti-β-actin (AC-15; Sigma). After incubation with HRP-labeled secondary antibodies, proteins were visualized using Western Lightning Plus-ECL (PerkinElmer) and an Amersham Imager 600 imaging system (GE Healthcare).

### Viral infectivity assay

Viral infectivity was assessed using TZM-bl cells (NIH AIDS Research & Reference Program), as described previously (25). The cells (1 × 10^5^ cells/well) were seeded onto 24-well plates and infected with serially diluted viruses normalized to the concentration of p24 proteins. At 24 h post-infection, firefly luciferase activity was measured using an LB9507 luminometer (Berthold, Germany).

### Immunofluorescence

293T cells (3×10^4^ cells/well) were cultured in a 2-well chamber slide and transfected with or without 100 ng pRad18-Myc. Cells were fixed in 3.6% formaldehyde in PBS and permeabilized with 0.1% NP-40 in PBS at room temperature. The cells were incubated with anti-HIV-1 p24 (65-005; Bioacademia, Japan), anti-HA (HA-7), anti-HIV-1 Env (KD-247), anti-RAD18 antibody (ab188235 and ab79763; Abcam), Monoclonal anti-splicing factor SC-35 (SC-35, Sigma), anti-Coilin antibody (ab11822; Abcam), anti-PML (PG-M3; Santa Cruz), anti-SFPQ (PSF) mAb (RN014MW; MBL), purified mouse anti-SMN (610646; BD Transduction Lab), anti-Nucleolin mAb (4E2; MBL), anti-Nucleolin antibody (ab22758; Abcam), anti-FLAG antibody (M2; Sigma), and/or anti-RAD18 antibody (ab188235 and ab79763; Abcam) at a 1:300 dilution in PBS containing 3% BSA. Cells were then stained with donkey anti-rabbit IgG (A21206), Alexa Fluor 488, goat anti-Human IgG (A11013), Alexa Fluor 488, donkey anti-mouse IgG (A21203), Alexa Fluor 594, donkey anti-rabbit IgG antibody, Alexa Fluor 647 (A315719), anti-Myc-tag mAb-Alexa Fluor 594 (M047-A59; MBL), and/or anti-Myc-tag mAb-Alexa Fluor488 (M047-A48; MBL) at a 1:300 dilution in PBS containing 3% BSA. The nuclei were stained with DAPI. Following extensive washing with PBS, the cells were mounted on slides using a mounting media of SlowFade Gold anti-fade reagent (Invitrogen). Samples were viewed using a confocal laser-scanning microscope (FV1200; Olympus, Tokyo, Japan; Zeiss LSM980 with Airyscan 2, Carl Zeiss, Oberkochen, Germany).

### Immunoprecipitation

Cells were lysed in a buffer containing 50 mM Tris-HCl (pH 8.0), 150 mM NaCl, 4 mM EDTA, 0.1% NP-40, 10 mM NaF, and a protease inhibitor cocktail (Nakalai Tesque). Lysates were pre-cleared with 30 µL of protein-G-Sepharose (GE). Pre-cleared supernatants were incubated with 5 µg of anti-Myc-tag mAb (PL-14; MBL) at 4°C for 1h. Following the absorption reaction of the precipitates with 30 µL of protein-G-Sepharose resin for 1h, the resin was washed four times with 700 µL of lysis buffer. Proteins were eluted by boiling the resin for 5 min in 2X Laemmli sample buffer. The proteins were then subjected to SDS-PAGE, followed by immunoblotting analysis using anti-Myc-tag mAb (PL-14; MBL), anti-FLAG (M2; Sigma), anti-HIV-1 GP41 (20-HR92; Fitzgerald), anti-KD247 (66), Anti-HIV-1 Rev (10; Icosagen), or Living Colors A.v. monoclonal antibody (JL-8; Clontech).

## Acknowledgements

We are grateful to Drs. Didier Trono and Priscilla Turelli (Ecole Polytechnique Fédérale de Lausanne, Switzerland) for their valuable suggestions. We would like to thank Dr. Didier Trono, Akio Adachi, Yoshio Koyanagi, and Yosuke Maeda for HIV-1 molecular clones and HIV-1 Env expression vector. This work was supported by the Japan Society for the Promotion of Science (JSPS) KAKENHI (Grant Number JP25065254) and the Japan Agency for Medical Research and Development (AMED) (Grant Number JP25fm0208101). K.S. and E.M. are supported by the Intramural Research Program of the NIH.

